# ACE: Explaining cluster from an adversarial perspective

**DOI:** 10.1101/2021.02.08.428881

**Authors:** Yang Young Lu, Timothy C. Yu, Giancarlo Bonora, William Stafford Noble

## Abstract

A common workflow in single-cell RNA-seq analysis is to project the data to a latent space, cluster the cells in that space, and identify sets of marker genes that explain the differences among the discovered clusters. A primary drawback to this three-step procedure is that each step is carried out independently, thereby neglecting the effects of the nonlinear embedding and inter-gene dependencies on the selection of marker genes. Here we propose an integrated deep learning framework, Adversarial Clustering Explanation (ACE), that bundles all three steps into a single workflow. The method thus moves away from the notion of “marker genes” to instead identify a panel of explanatory genes. This panel may include genes that are not only enriched but also depleted relative to other cell types, as well as genes that exhibit differences between closely related cell types. Empirically, we demonstrate that ACE is able to identify gene panels that are both highly discriminative and nonredundant, and we demonstrate the applicability of ACE to an image recognition task. ^1^

## 1. Introduction

Single-cell sequencing technology has enabled the high-throughput interrogation of many aspects of genome biology, including gene expression, DNA methylation, histone modification, chromatin accessibility and genome 3D architecture (Stuart & Satija, 2019) In each of these cases, the resulting high-dimensional data can be represented as a sparse matrix in which rows correspond to cells and columns correspond to features of those cells (gene expression values, methylation events, etc.). Empirical evidence suggests that this data resides on a low-dimensional manifold with latent semantic structure (Welch et al., 2017). Accordingly, identifying groups of cells in terms of their inherent latent semantics and thereafter reasoning about the differences between these groups is an important area of research (Plumb et al., 2020).

In this study, we focus on the analysis of single cell RNA-seq (scRNA-seq) data. This is the most widely available type of single-cell sequencing data, and its analysis is challenging not only because of the data’s high dimensionality but also due to noise, batch effects, and sparsity (Amodio et al., 2019). The scRNA-seq data itself is represented as a sparse, cell-by-gene matrix, typically with tens to hundreds of thousands of cells and tens of thousands of genes. A common workflow in scRNA-seq analysis (Pliner et al., 2019) consists of three steps: (1) learn a compact representation of the data by projecting the cells to a lower-dimensional space; (2) identify groups of cells that are similar to each other in the low-dimensional representation, typically via clustering; and (3) characterize the differences in gene expression among the groups, with the goal of understanding what biological processes are relevant to each group. Optionally, known “marker genes” may be used to assign cell type labels to the identified cell groups.

A primary drawback to the above three-step procedure is that each step is carried out independently. Here, we propose an integrated, deep learning framework for scRNA-seq analysis, Adversarial Clustering Explanation (ACE), that projects scRNA-seq data to a latent space, clusters the cells in that space, and identifies sets of genes that succinctly explain the differences among the discovered clusters (Figure 1). At a high level, ACE first “neuralizes” the clustering procedure by reformulating it as a functionally equivalent multi-layer neural network (Kauffmann et al., 2019). In this way, in concatenation with a deep autoencoder that generates the low-dimensional representation, ACE is able to attribute the cell’s group assignments all the way back to the input genes by leveraging gradient-based neural network explanation methods. Next, for each sample, ACE seeks small perturbations of its input gene expression profile that lead the neuralized clustering model to alter the group assignments. These adversarial perturbations allow ACE to define a concise gene set signature for each cluster or pair of clusters. In particular, ACE attempts to answer the question, “For a given cell cluster, can we identify a subset of genes whose expression profiles are sufficient to identify members of this cluster?” We frame this problem as a ranking task, where thresholding the ranked list yields a set of explanatory genes.

**Figure 1.**
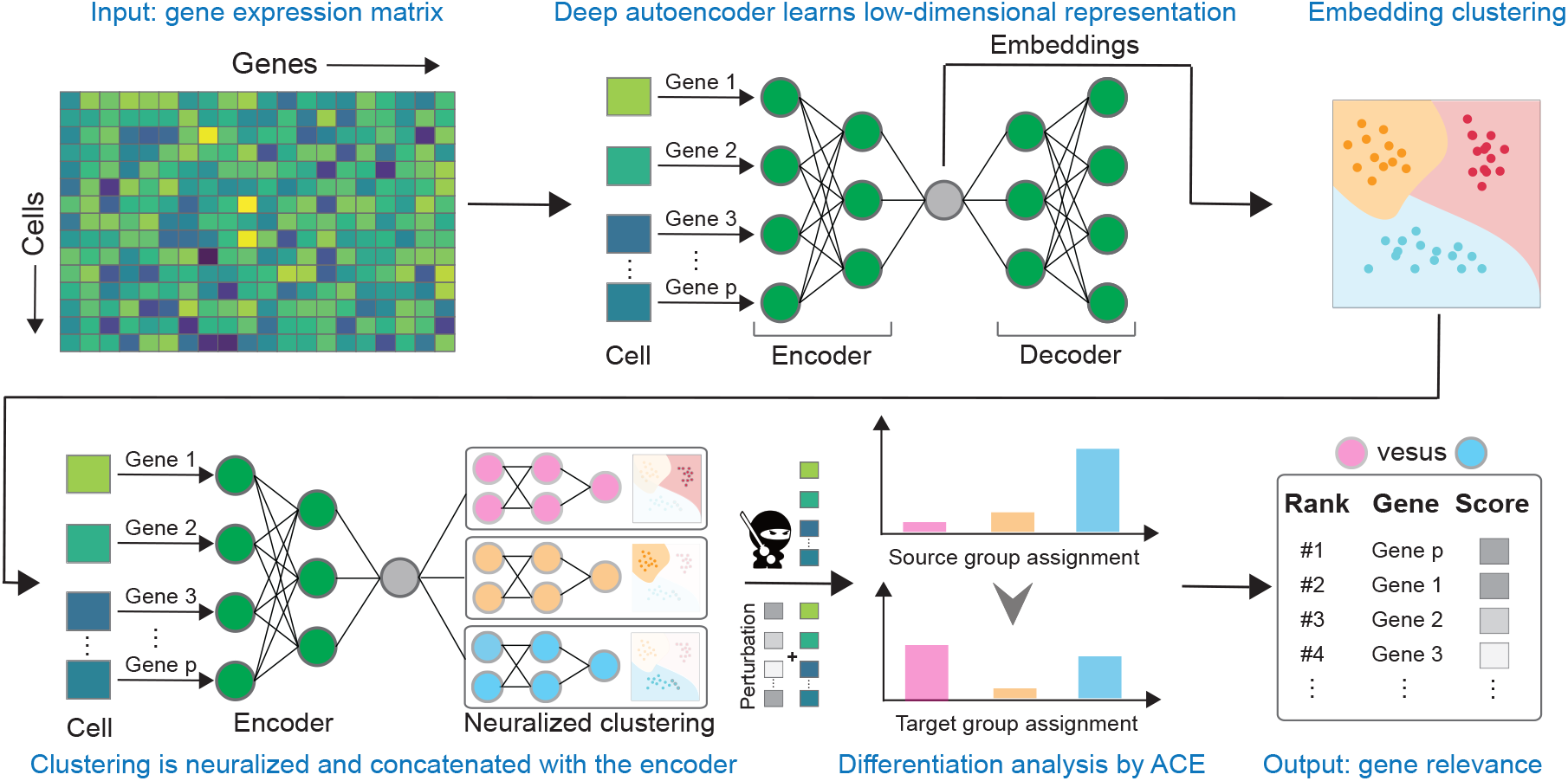
ACE workflow. ACE takes as input a single-cell gene expression matrix and learns a low-dimensional representation for each cell. Next, a neuralized version of the *k*-means algorithm is applied to the learned representation to identify cell groups. Finally, for pairs of groups of interest (either each group compared to its complement, or all pairs of groups), ACE seeks small perturbations of its input gene expression profile that lead the neuralized clustering model to alter the assignment from one group to the other. The workflow employs a combined objective function to induce the nonlinear embedding and clustering jointly. ACE produces as output the learned embedding, the cell group assignments, and a ranked list of explanatory genes for each cell group.

ACE’s joint modeling approach offers several benefits relative to the existing state of the art. First, most existing methods for the third step of the analysis pipeline—identifying genes associated with a given group of cells—treat each gene independently (Love et al., 2014). These approaches ignore the dependencies among genes that are induced by gene networks, and often yield lists of genes that are highly redundant. ACE, in contrast, aims to find a small set of genes that jointly explain a given cluster or pair of clusters. Second, most current methods identify genes associated with a group of cells without considering the nonlinear embedding model which maps the gene expression to the low-dimensional representation where the groups are defined in the first place. To our knowledge, the only exception is the global counterfactual explanation (GCE) algorithm (Plumb et al., 2020) which is motivated by compressed sensing (Candès, 2006). A third advantage of ACE’s integrated approach is its ability to take into account batch effects during the assignment of genes to clusters. Standard nonlinear embedding methods, such as t-SNE (Van der Maaten & Hinton, 2008) and UMAP (McInnes & Healy, 2018; Becht et al., 2019), cannot take such structure into account and hence may lead to incorrect interpretation of the data (Amodio et al., 2019; Li et al., 2020). To address this problem, deep autoencoders with integrated denoising and batch correction can be used for scRNA-seq analysis (Lopez et al., 2018; Amodio et al., 2019; Li et al., 2020). We demonstrate below that batch effect structure can be usefully incorporated into the ACE model.

A notable feature of ACE’s approach is that, by identifying genes jointly, the method moves away from the notion of a “marker gene” to instead identify a “gene panel”. As such, genes in the panel may not be solely enriched in a single cluster, but may together be predictive of the cluster. In particular, in addition to a ranking of genes, ACE assigns a Boolean to each gene indicating whether its inclusion in the panel is positive or negative, i.e., whether the gene’s expression is enriched or depleted relative to cluster membership. We have applied ACE to both simulated and real datasets to demonstrate its empirical utility. Our experiments demonstrate that ACE identifies gene panels that are highly discriminative and exhibit low redundancy. We further provide results suggesting that ACE is useful in domains beyond biology, such as image recognition.

## 2. Related work

ACE falls into the paradigm of deep neural network interpretation methods, which have been developed primarily in the context of classification problems. These methods can be loosely categorized into three types: feature attribution methods, counterfactual-based methods, and model-agnostic approximation methods. Feature attribution methods assign an importance score to individual features so that higher scores indicate higher importance to the output prediction (Simonyan et al., 2013; Shrikumar et al., 2017; Lundberg & Lee, 2017). Counterfactual-based methods typically identify the important subregions within an input sample by perturbing the subregions (by adding noise, rescaling (Sundararajan et al., 2017), blurring (Fong & Vedaldi, 2017), or inpainting (Chang et al., 2018)) and measuring the resulting changes in the predictions. Lastly, model-agnostic approximation methods approximate the model being explained by using a simpler, surrogate function which is self-explainable (*e.g.,* a sparse linear model, etc.) (Ribeiro et al., 2016). Recently, some interpretation methods have emerged to understand models beyond classification tasks (Samek et al., 2020; Kauffmann et al., 2020; 2019), including the one we present in this paper for the purpose of cluster explanation.

ACE’s perturbation approach draws inspiration from adversarial machine learning (Xu et al., 2020) where imperceivable perturbations are maliciously crafted to mislead a machine learning model to predict incorrect outputs. In particular, ACE’s approach is closest to the setting of a “white-box attack,” which assumes complete knowledge to the model, including its parameters, architecture, gradients, etc. (Szegedy et al., 2013; Kurakin et al., 2016; Madry et al., 2017; Carlini & Wagner, 2017). In contrast to these methods, ACE re-purposes the malicious adversarial attack for a constructive purpose, identifying sets of genes that explain clusters in scRNA-seq data.

ACE operates in concatenation with a deep autoencoder that generates the low-dimensional representation. In this paper, ACE uses SAUCIE (Amodio et al., 2019), a commonly-used scRNA-seq embedding method that incorporates batch correction. In principle, ACE is generalizable to any off-the-shelf scRNA-seq embedding methods, including SLICER (Welch et al., 2016), scVI (Lopez et al., 2018), scANVI (Xu et al., 2021), DESC (Li et al., 2020), and ItClust (Hu et al., 2020).

## 3. Approach

### 3.1. Problem setup

We aim to carry out three analysis steps for a given scRNA-seq dataset, producing a low-dimensional representation of each cell’s expression profile, a cluster assignment for each cell, and a concise set of “explanatory genes” for each cluster or pair of clusters. Let *X* = (*x*_1_, *x*_2_, … , *x*_*n*_)^*T*^ ∈ ℝ^*n×p*^ be the normalized gene expression matrix obtained from a scRNA-seq experiment, where rows correspond to *n* cells and columns correspond to *p* genes. ACE relies on the following three components: (1) an autoencoder to learn a low-dimensional representation of the scRNA-seq data, (2) a neuralized clustering algorithm to identify groups of cells in the low-dimensional representation, and (3) an adversarial perturbation scheme to explain differences between groups by identifying explanatory gene sets.

### 3.2. Learning the low-dimensional representation

Embedding scRNA-seq expression data into a low-dimensional space aims to capture the underlying structure of the data, based upon the assumption that the biological manifold on which cellular expression profiles lie is inherently low-dimensional. Specifically, ACE aims to learn a mapping *f* (·): ℝ^*p*^ ↦ ℝ^*d*^ that transforms the cells from the high-dimensional input space ℝ^*p*^ to a lower-dimensional embedding space ℝ^*d*^, where *d* ≪ *p*. To accurately represent the data in ℝ^*d*^, we use an autoencoder consisting of two components, an encoder *f* (·): ℝ^*p*^ ↦ ℝ^*d*^ and a decoder *g*(·): ℝ^*d*^ ↦ ℝ^*p*^. This autoencoder optimizes the generic loss

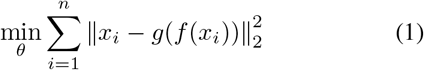

Finally, we denote *Z* = (*z*_1_, *z*_2_, … ,*z*_*n*_) ∈ ℝ^*n×d*^ as the low-dimensional representation obtained from the encoder, where *z*_*i*_ ∈ ℝ^*d*^ = *f*(*x*_*i*_) is the embedded representation of cell *x*_*i*_.

The autoencoder in ACE can be extended in several important ways. For example, in some settings, Equation 1 is augmented with a task-specific regularizer Ω(*X*):

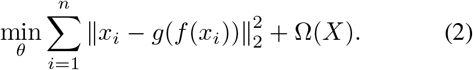

As mentioned in Section 2, the scRNA-seq embedding method used by ACE, SAUCIE, encodes in Ω(*X*) a batch correction regularizer by using maximum mean discrepancy. In this paper, ACE uses SAUCIE coupled with a feature selection layer (Abid et al., 2019), with the aim of minimizing redundancy and facilitating selection of diverse explanatory gene sets.

### 3.3. Neuralizing the clustering step

To carry out clustering in the low-dimensional space learned by the autoencoder, ACE uses a neuralized version of the *k*-means algorithm. This clustering step aims to partition *Z* ∈ ℝ^*n×d*^ into *C* groups, where each group potentially corresponds to a distinct cell type.

The standard *k*-means algorithm aims to minimize the following objective function by identifying a set of group centroids {*μ*_*c*_ ∈ ℝ^*d*^: *c* = 1, 2, … , *C*}:

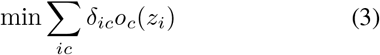

where *δ*_*ic*_ indicates whether cell *z*_*i*_ belongs to group *c* and the “outlierness” measure *o*_*c*_ (*z*_*i*_) of cell *z*_*i*_ relative to group *c* is defined as *o*_*c*_ (*z*_*i*_) = ||*z*_*i*_ − *μ*_*c*_||^2^.

Following Kauffmann et al. (2019), we neuralize the *k*-means algorithm by creating a neural network containing *C* modules, each with two layers. The architecture is motivated by a soft assignment function that quantifies, for a particular cell *z*_*i*_ and a specified group *c*, the group assignment probability score

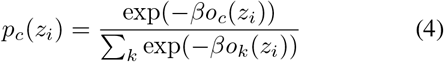

where the hyperparameter *β* controls the clustering fuzziness. As *β* approaches infinity, Equation 4 approaches the indicator function for the closest centroid and thus reduces to hard clustering. To measure the confidence of group assignment, we use a logit function written as

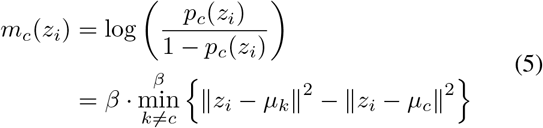

where 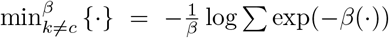 indicates a soft min-pooling layer. (See Kauffmann et al. (2019) for a detailed derivation.) The rationale for using the logit function is that if there is as much confidence <u>supporting</u> the group membership as <u>against</u> it, then the confidence score *m*_*c*_(*z*) = 0. Additionally, Equation 5 has the following interpretation: the data point *z* belongs to the group *c* if and only if the distance to its centroid is smaller than the distance to all other competing groups. Equation 5 further decomposes into a two-layer neural network module:

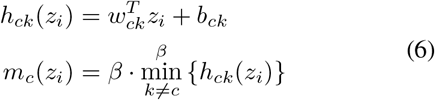

where the first layer is a linear transformation layer with parameters *w*_*ck*_ = 2 · (*μ*_*c*_ – *μ*_*k*_) and *b*_*ck*_ = ||*μ*_*k*_||^2^ – ||*μ*_*c*_||^2^, and the second layer is the soft min-pooling layer introduced in Equation 5. ACE constructs one such module for each of the *C* clusters, as illustrated in Figure 1.

### 3.4. Explaining the groups

ACE’s final step aims to induce, for each cluster identified by the neuralized *k*-mean algorithm, a ranking on genes such that highly ranked genes best explain that cluster. We consider two variants of this task: the <u>one-vs-rest</u> setting compares the group of interest *Z*_*s*_ = *f* (*X*_*s*_) ⊆ *Z* to its complement set *Z*_*t*_ = *f* (*X*_*t*_) ⊆ *Z*, where *X*_*t*_ = *X*\*X*_*s*_; the <u>one-vs-one</u> setting compares one group of interest in *Z*_*s*_ = *f* (*X*_*s*_) ⊆ *Z* to a second group of interest *Z*_*t*_ = *f*(*X*_*t*_) ⊆ *Z*. In each setting, the goal is to identify the key differences between the source group *X*_*s*_ ⊆ *X* and the target group *X*_*t*_ ⊆ *X* in the input space, *i.e.,* in terms of the genes.

We treat this as a neural network explanation problem by finding the minimal perturbation within the group of interest, *x* ∈ *X*_*s*_, that alters the group assignment from the source group *s* to the target group *t*. Specifically, we optimize an objective function that is a mixture of two terms: the first term is the difference between the current sample *x* and the perturbed sample 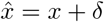 where *δ* ∈ ℝ^*p*^, and the second term quantifies the difference in group assignments induced by the perturbation.

The objective function for the one-vs-one setting is

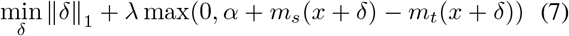

where *λ* > 0 is a tradeoff coefficient to either encourage a small perturbation of *x* when small or a stronger alternation to the target group when large. The second term penalizes the situation where the group logit for the source group *s* is still larger than the target group *t*, up to a pre-specified margin *α* > 0. In this paper we fix *α* = 1.0. The difference between the current sample *x* and the potentially perturbed 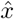 is measured by the *L*_1_ norm to encourage sparsity and non-redundancy. Note that Equation 7 assumes that the input expression matrix is normalized so that a perturbation added to one gene is equivalent to that same perturbation added to a different gene.

Analogously, in the one-vs-rest case, the objective function for the optimization is

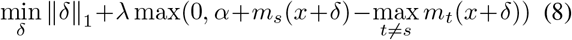

where the second term penalizes the situation in which the group logit for the source group *s* is larger than all non-source target groups.

Finally, with the *δ* ∈ ℝ^*p*^ obtained by optimizing either Equation 7 or Equation 8, ACE quantifies the importance of the *i*th gene relative to a perturbation from source group *s* to target group *t* as the absolute value of *δ*_*i*_, thereby inducing a ranking in which highly ranked genes are more specific to the group of interest.

## 4. Baseline methods

We compare ACE against six methodologically distinct baseline methods, each of which induces a ranking on genes in terms of group-specific importance, analogous to ACE. DESeq2 (Love et al., 2014) is a representative statistical hypothesis testing method that tests for differential gene expression based on a negative binomial model. The main caveat of DESeq2 is that it treats each gene as independent.

The Jensen-Shannon Distance (JSD) (Cabili et al., 2011) is a representative distribution distance-based method which quantifies the specificity of a gene to a cell group. Similar to DESeq2, JSD considers each gene independently.

Global counterfactual explanation (GCE) (Plumb et al., 2020) is a compressed sensing method that aims to identify consistent differences among all pairs of groups. Unlike ACE, GCE is mainly designed for the one-vs-one setting because it relies on an objective function that characterizes each group via the cluster centroid.

The gene relevance score (GRS) (Angerer et al., 2020) is a gradient-based explanation method that aims to attribute a low-dimensional embedding back to the genes. The main limitations of GRS are two-fold. First, the embedding used in GRS is constrained to be a diffusion map, which is chosen specifically to make the gradient easy to calculate. Second, taking the gradient with respect to the embedding only indirectly measures the group differentiation compared to taking the gradient with respect to the group difference directly, as in ACE.

SmoothGrad (Smilkov et al., 2017) and SHAP (Lundberg & Lee, 2017), which are designed primarily for classification problems, are two representative feature attribution methods. Each one computes an importance score that indicates each gene’s contribution to the clustering assignment. Smooth-Grad relies on knowledge to the model, whereas SHAP does not.

## 5. Results

### 5.1. Performance on simulated data

To compare ACE to each of the baseline methods, we used a recently reported simulation method, SymSim (Zhang et al., 2019), to generate two synthetic scRNA-seq datasets: one “clean” dataset and one “complex” dataset. In both cases, we simulated many redundant genes, in order to adequately challenge methods that aim to detect a minimal set of informative genes.

The simulation of the clean dataset uses a protocol similar to that of Plumb et al. (2020). We first used SymSim to generate a background matrix containing simulated counts from 500 cells, 2000 genes, and five distinct clusters. We then used this background matrix to construct our simulated dataset of 500 cells by 220 genes. The simulated data is comprised of three sets of genes: 20 causal genes, 100 dependent genes, and 100 noise genes. To select the causal genes, we identified all genes that are differentially expressed by SymSim’s criteria (nDiff-EVFgene > 0 and |log_2_ fold-change| > 0.8) between at least one pair of clusters, and we selected the 20 genes that exhibit the largest average fold-change across all pairs of clusters in which the gene was differentially expressed. A UMAP embedding on these causal genes alone confirms that they are jointly capable of separating cells into their respective clusters (Fig. 2A). Next, we simulated 100 dependent genes, which are weighted sums of 1–10 randomly selected causal genes, with added gaussian noise. As such, a dependent gene is highly correlated with a causal gene or with a linear combination of multiple causal genes. The weights were sampled from a continuous uniform distribution, *U* (0.01, 0.8), and the gaussian noise was sampled from *N* (0, 1). As expected, the dependent genes are also jointly capable of separating cells into their respective clusters (Fig. 2A). Lastly, we found all genes that were not differentially expressed between any cluster pair in the ground truth, and we randomly sampled 100 noise genes. These genes provide no explanation of the clustering structure (Fig. 2A).

**Figure 2.**
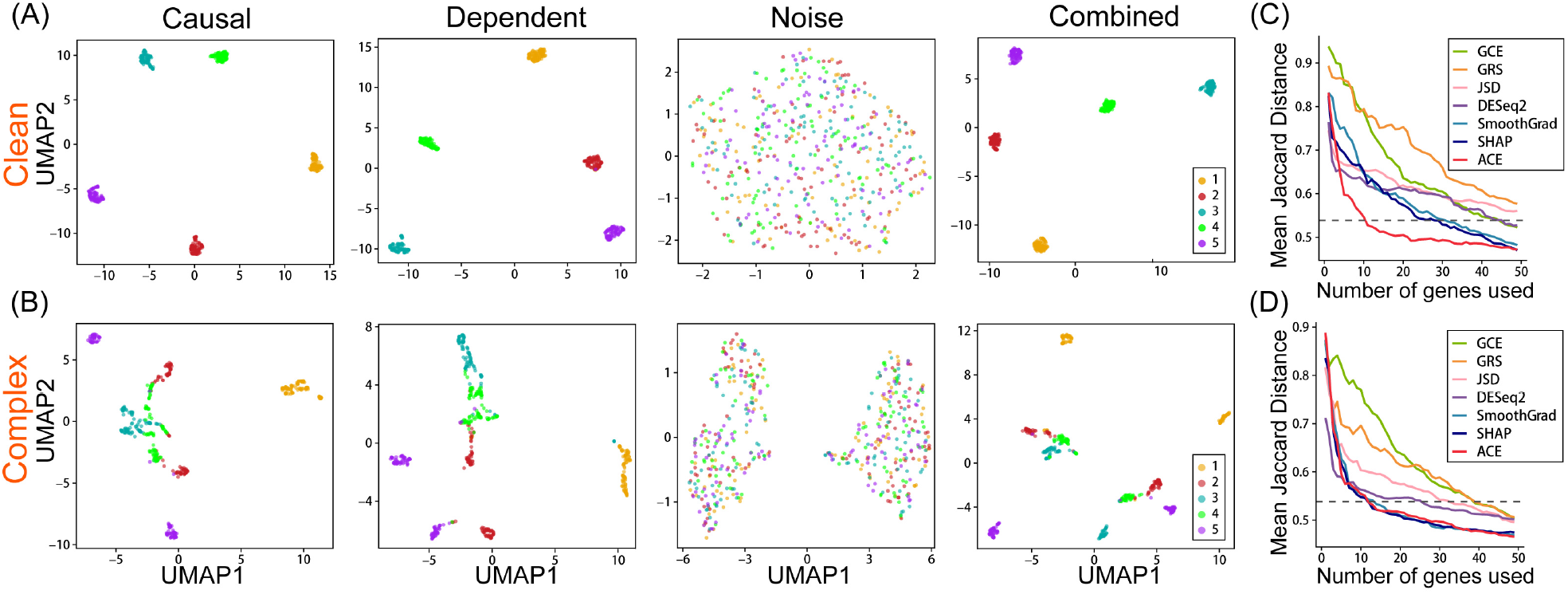
Comparing ACE to baseline methods on simulated scRNA-seq datasets. Each dataset consists of 20 causal genes, 100 dependent genes, and 100 noise genes. (**A**) UMAP embeddings of cells composing the clean dataset. Panels correspond to embeddings using the three subsets of genes (causal, dependent, and noise), as well as all of the genes together. (**B**) Same as panel A, but for the complex dataset. (**C**) Comparison of methods via Jaccard distance as a function of the number of genes in the ranking. ACE performs substantially better than each of the baseline methods on the clean dataset. The gray dashed line indicates the mean Jaccard distance achieved by the 20 causal genes alone. (**D**) Same as panel C but for the complex dataset.

To simulate the complex dataset, we used SymSim to add dropout events and batch effects to the background matrix generated previously. We then selected the same exact causal and noise genes as in the clean dataset, and used the same exact random combinations and weights to generate the dependent genes. Thus, the clean and complex datasets contain the same 220 genes; however, the complex dataset enables us to gauge how robust ACE is to artifacts of technical noise observed in real single-cell RNA-seq datasets (Fig. 2B).

To compare the different gene ranking methods, we need to specify the ground truth cluster labels and a performance measure. We observe that the embedding representation learned by ACE exhibits clear cluster patterns even in the presence of dropout events and batch effects, and thus ACE’s *k*-means clustering is able to recover these clusters (Appendix Figure A.1). Accordingly, to compare different methods for inducing gene rankings, we provide ACE and each baseline method with the ground truth clustering labels from the original study (Zheng et al., 2017). ACE then calculates the group centroid used in Equation 3 by averaging the data points of the corresponding ground truth cluster. The embedding layer together with the group centroids are then used to build the neuralized clustering model (Equation 6). Each method produces gene rankings for every cluster in a one-vs-rest fashion. To measure how well a gene ranking captures clustering structure, we use the Jaccard distance to measure the similarity between a cell’s *k* nearest neighbors (*k*-NN) when using a subset of top-ranked genes and a cell’s *k*-NN when using all genes. To compute the *k*-NN, we use the Euclidean distance metric. The Jaccard distance is defined as

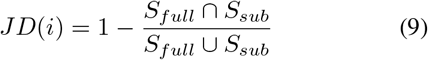

where *S*_*full*_ represents cell *i*’s *k*-NN’s when using all genes, and *S*_*sub*_ represents cell *i*’s *k*-NN’s when using a subset of top-ranked genes. If the subset of top-ranked genes does a good job of explaining a cluster of cells, then *S*_*full*_ ∩ *S*_*sub*_ and *S*_*full*_ ∪ *S*_*sub*_ should be nearly equal, and the Jaccard distance should approach 0. We select the gene ranking used to derive a subset of top-ranked genes based on the cell cluster assignment. For example, if the cell belongs in cluster 2, we use the cluster 2 vs. rest gene ranking. Thus, to obtain a global measure of how well a clustering structure is captured on a subset of top-ranked genes, we report the mean Jaccard distance across all cells.

Our analysis shows that ACE considerably outperforms each of the baseline methods on the clean dataset, indicating that it is superior at identifying the minimal set of informative genes (Fig. 2B). Notably, ACE outperforms the mean Jaccard distance achieved by the causal genes alone before reaching 20 genes used, suggesting that the method successfully identifies dependent genes that are more informative than individual causal genes. ACE also performs strongly on the complex dataset, though it appears to perform on par with SmoothGrad and SHAP) (Fig. 2D). Notably, these three methods —ACE, SHAP, and SmoothGrad —share a common feature, employing the SAUCIE framework that facilitates automatic batch effect correction, highlighting the utility of DNN-based dimensionality reduction and interpretation methods for single-cell RNA-seq applications.

### 5.2. Real data analysis

We next applied ACE to a real dataset of peripheral blood mononuclear cells (PBMCs) (Zheng et al., 2017), represented as a cell-by-gene log-normalized expression matrix containing 2638 cells and 1838 highly variable genes. The cells in the dataset were previously categorized into eight cell types, obtained by performing Louvain clustering (Blondel et al., 2008) and annotating each cluster on the basis of differentially expressed marker genes. As shown in Figure 3A and Appendix Figure A.2, ACE’s *k*-means clustering successfully recovers the reported cell types based upon the 10-dimensional embedding learned by SAUCIE.

**Figure 3.**
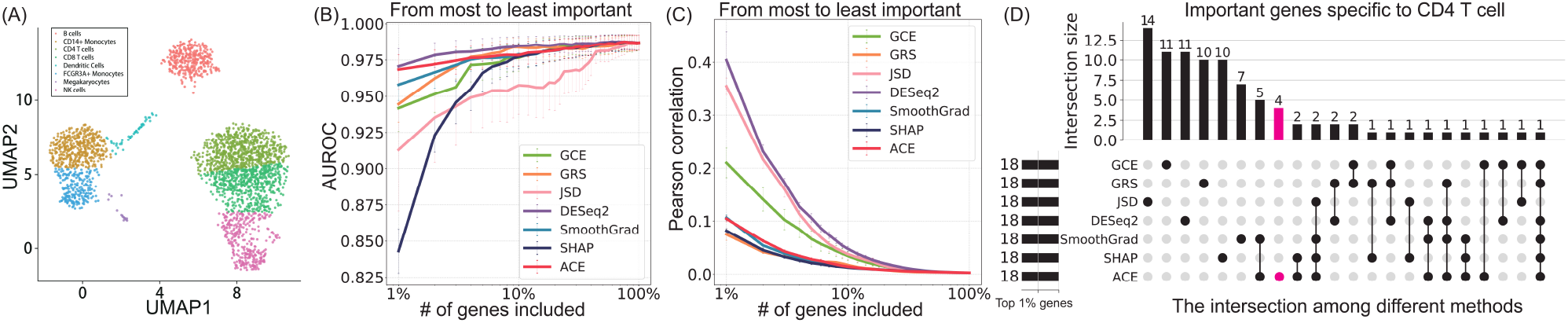
Comparing ACE to baseline methods on PBMC dataset. (**A**) UMAP embedding of PBMC cells labelled by ACE’s *k*-means clustering assignment. (**B**) Classification performance of each method, as measured by AUROC, as a function of the number of genes in the set. Error bars correspond to the standard error of the mean of AUROC scores from each test split across different target groups. (**C**) Redundancy among the top *k* genes, as measured by Pearson correlation, as a function of *k*. Error bars correspond to the standard error of the mean calculated from the group-specific correlations. (**D**) The figure plots overlaps among the top 18 genes (corresponding to 1% of 1838 genes) identified by all seven methods with respect to the CD4 T cell cluster.

We first aimed to quantify the discriminative power of the top-ranked genes identified by ACE in comparison to the six baseline methods. To do this, we applied all the six baseline methods to the PBMC dataset using the groups identified by the *k*-means clustering based on the SAUCIE embedding. For each group of cells, we extracted the top-*k* group-specific genes reported by each method, where *k* ranges from 1%, 2%, … ,100% among all genes. Given the selected gene subset, we then trained a support vector machine (SVM) classifier with a radial basis function kernel to separate the target group from the remaining groups. The SVM training involves two hyperparameters, the regularization coefficient *C* and the bandwidth parameter *σ*. The *σ* parameter is adaptively chosen so that the training data is Z-score normalized, using the default settings in Scikit-learn (Pedregosa et al., 2011). The *C* parameter is selected by grid search from 5^−5^, 5^−4^, … , 5^0^, … , 5^4^, 5^5^}. The classification performance, in terms of area under the receiver operating characteristic curve (AUROC), is evaluated by 3-fold stratified cross-validation, and an additional 3-fold cross-validation is applied within each training split to determine the optimal *C* hyperparameter. Finally, AUROC scores from each test split across different target groups are aggregated and reported, in terms of the mean and the standard error of the mean. Two cell types—megakaryocytes and dendritic cells—are excluded due to insufficient sample size (< 50). As shown in Figure 3B, the top-ranked genes reported by ACE are among the most discriminative across all methods, particularly when the inclusion size is small (≤3%). The only method that yields superior performance is DESeq2.

We next tested the redundancy of top-ranked genes, as it is desirable to identify diverse explanatory gene sets with minimum redundancy. Specifically, for each target group of cells, we calculate the Pearson correlations between all gene pairs within top *k* genes, for varying values of *k*. The mean and standard error of the mean of these correlations are computed within each group and then averaged across different target groups. The results of this analysis (Figure 3C) suggest that the top-ranked genes reported by ACE are among the least redundant across all methods. Other methods that exhibit low redundancy include GRS and the two methods that use the same SAUCIE model (*i.e.,* SmoothGrad and SHAP). In conjunction with the discriminative power analysis in Figure 3B, we conclude that ACE achieves a powerful combination of high discriminative power and low redundancy.

Finally, to better understand how these methods differ from one another, we investigated the consistency among the top-ranked genes reported by each method. For this analysis, we focused on one particular group, CD4 T cells. We discover strong disagreement among the methods (Figure 3D). Surprisingly, no single gene is selected among the top 1% by all methods. Among all methods, ACE covers the most that are reported by at least one other method (14 out of 18 genes). The four genes that ACE uniquely identifies (red bar in Figure 3D)—CCL5, GZMK, SPOCD1, and SNRNP27—are depleted rather than enriched relative to other cell types. It is worth mentioning that both CCL5 and GZMK are enriched in CD8 T cells (Thul et al., 2017), the closest cell type to CD4 T cell (Figure 3A). This observation suggests ACE identifies cells that exhibit highly discriminative changes in expression between two closely related cell types. Indeed, among ACE’s 18-gene panel, 15 genes are depleted rather than enriched, suggesting that much of CD4’s cell identity may be due to inhibition rather than activation of specific genes. In summary, ACE is able to move away from the notion of a “marker gene” to instead identify a highly discriminative, nonredundant gene panel.

### 5.3. Image analysis

Although we developed ACE for application to scRNA-seq data, we hypothesized that the method would be useful in domains beyond biology. Explanation methods are potentially useful, for example, in the analysis of biomedical images, where the explanations can identify regions of the image responsible for assignment of the image to a particular phenotypic category. As a proof of principle for this general domain, we applied ACE to the MNIST handwritten digits dataset (LeCun, 1998), with the aim of studying whether ACE can identify which pixels in a given image explain why the image was assigned to one digit versus another. Specifically, we solve the optimization problem for each input image in Equation 7, seeking an image-specific set of pixel modifications, subject to the constraint that the perturbed image pixel values are restricted to lie in the range [0, 1].

Note that this task is somewhat different from the scRNA-seq case: in the MNIST case, ACE finds a different set of explanatory pixels for each image, whereas in the scRNA-seq case, ACE seeks a single set of genes that explains label differences across all cells in the dataset.

ACE was applied to this dataset as follows. We used a simple convolution neural network architecture containing two convolution layers, each with a modest filter size (5, 5), a modest number of filters (32) and ReLU activation, followed by a max pooling layer with a pool size (2, 2), a fully connected layer, and a softmax layer. The model was trained on the MNIST training set (60,000 examples) for 10 epochs, using Adam (Kingma & Ba, 2015) with an initial learning rate of 0.001. The network achieves 98.7% classification accuracy on the test test of 10,000 images. We observe that the embedding representation in the last pooling layer exhibits well-separated cluster patterns (Appendix Figure A.3). Since our goal is not to learn the cluster structure per se, for simplicity, we fixed the number of groups to be the number of digit categories (*i.e.,* 10) and calculated the group centroid used in Equation 3 by averaging the data points of the corresponding category. The embedding layer together with the group centroids are then used to build the neuralized clustering model (Equation 6.)

The results of this analysis show that ACE does a good job of identifying sets of pixels that accurately explain differences between pairs of digits. We examined the pixel-wise explanations of 20 pairs of digits, randomly selected to cover each digit category at least once in both directions (Fig. 4). For example, to convert “8” to “5,” ACE disconnects the top right and bottom left of “8,” as expected. Similarly, to convert “8” to “3,” ACE disconnects the top left and bottom left of “8.” It is worth noting that the modifications introduced by ACE are inherently symmetric. For example, to convert “1” to “7” and back again, ACE suggests adding and removing the same part of “7.”

**Figure 4.**
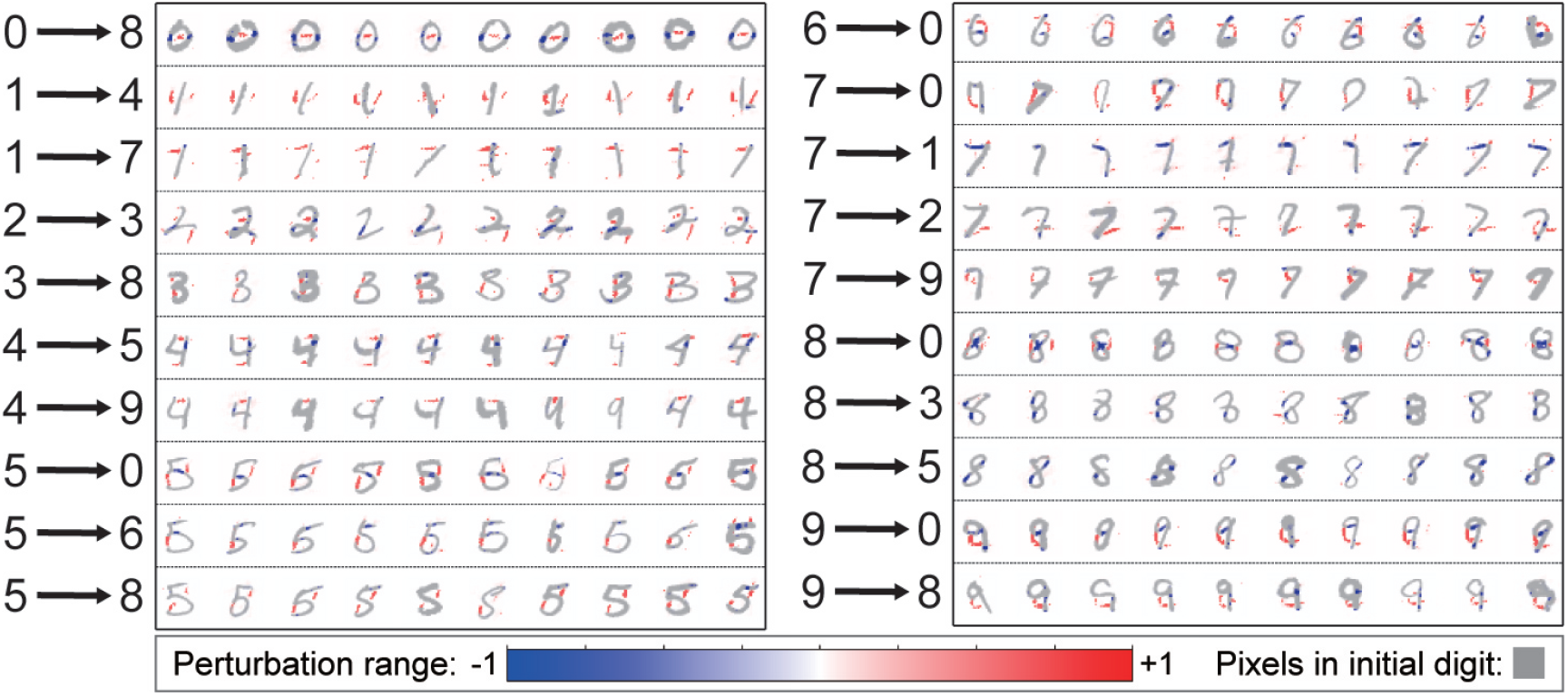
Applying ACE to the MNIST dataset. ACE is able to explain 20 types of digit transitions in a pixel-wise manner. These digit transitions are chosen such that each digit category is covered at least once in both directions.

## 6. Discussion and conclusion

In this work, we have proposed a deep learning-based scRNA-seq analysis pipeline, ACE, that projects scRNA-seq data to a latent space, clusters the cells in that space, and identifies sets of genes that succinctly explain the differences among the discovered clusters. Compared to existing state-of-the-art methods, ACE jointly takes into consideration both the nonlinear embedding of cells to a low-dimensional representation and the intrinsic dependencies among genes. As such, the method moves away from the notion of a “marker gene” to instead identify a panel of genes. This panel may include genes that are not only enriched but also depleted relative to other cell types, as well as genes that exhibit important differences between closely related cell types. Our experiments demonstrate that ACE identifies gene panels that are highly discriminative sets and exhibit low redundancy. We also provide results suggesting that ACE’s approach may be useful in domains beyond biology, such as image recognition.

This work points to several promising directions for future research. In principle, ACE can be used in conjunction with any off-the-shelf scRNA-seq embedding method. Thus, empirical investigation of the utility of generalizing ACE to use embedders other than SAUCIE would be interesting. Another possible extension is to apply neuralization to alternative clustering algorithms. For example, in the context of scRNA-seq analysis the Louvain algorithm (Blondel et al., 2008) is commonly used and may be a good candidate for neuralization. A promising direction for future work is to provide confidence estimation for the top-ranked group-specific genes, in terms of q-values (Storey, 2003), with the help of the recently proposed knockoffs framework (Barber & Candès, 2015; Lu et al., 2018).

---

**Figure A.1.**
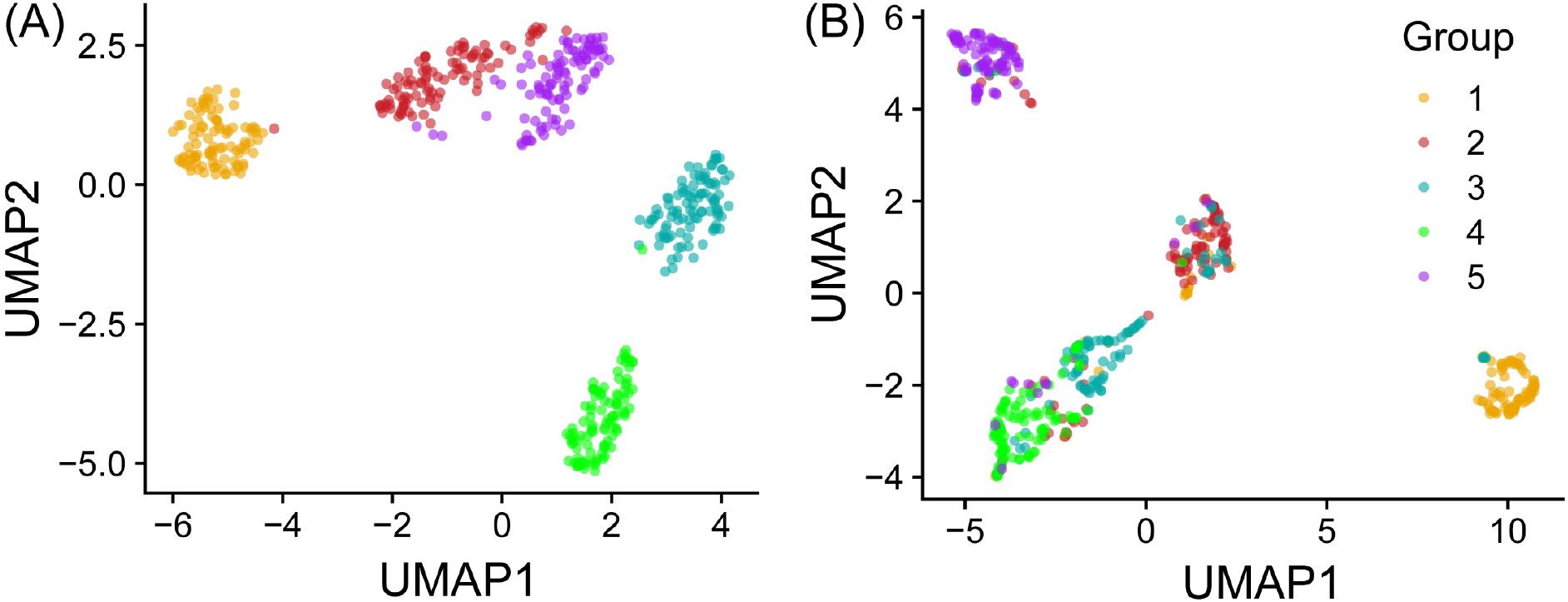
The embedding representation learned by SAUCIE exhibits well-separated cluster patterns on both (**A**) clean and (**B**) complex simulated scRNA-seq datasets.

**Figure A.2.**
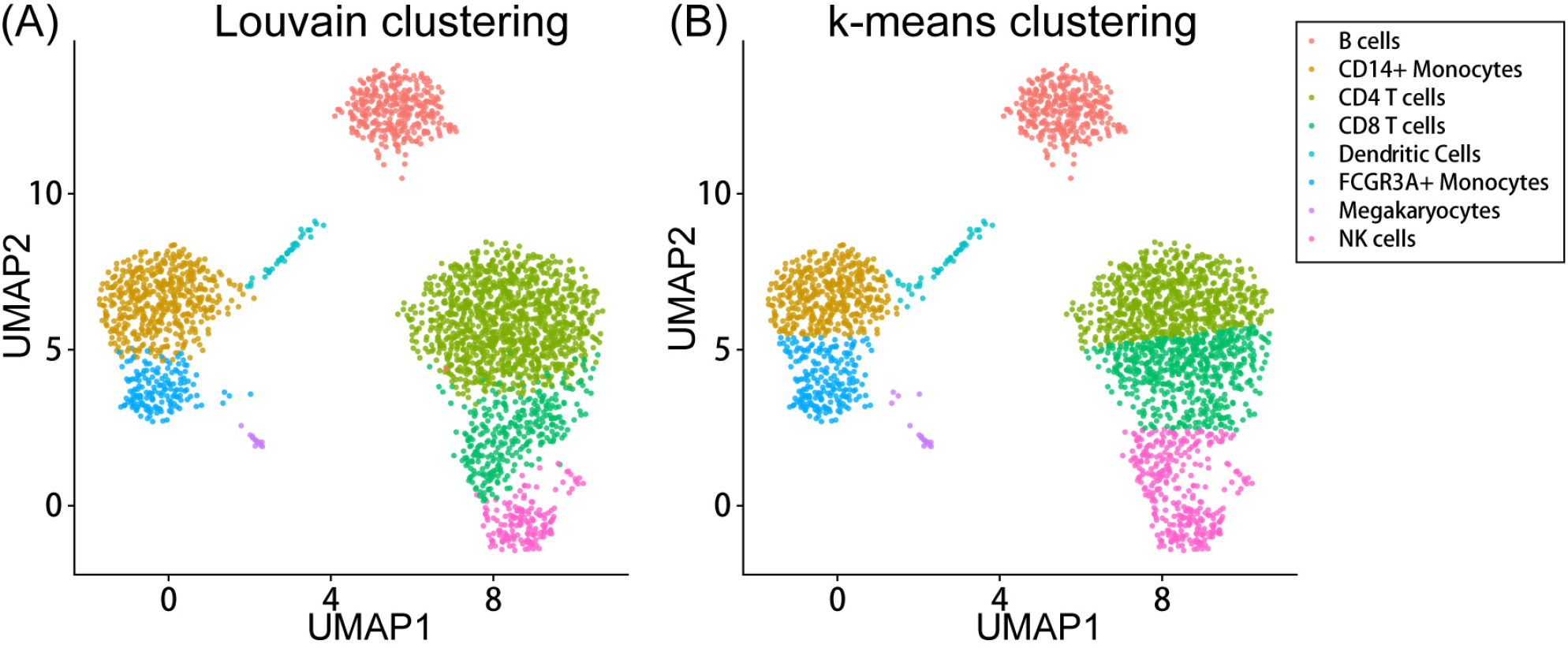
The embedding representation learned by SAUCIE exhibits similar cluster patterns by using either (**A**) the Louvain algorithm or (**B**) *k*-means clustering on the PBMC dataset.

**Figure A.3.**
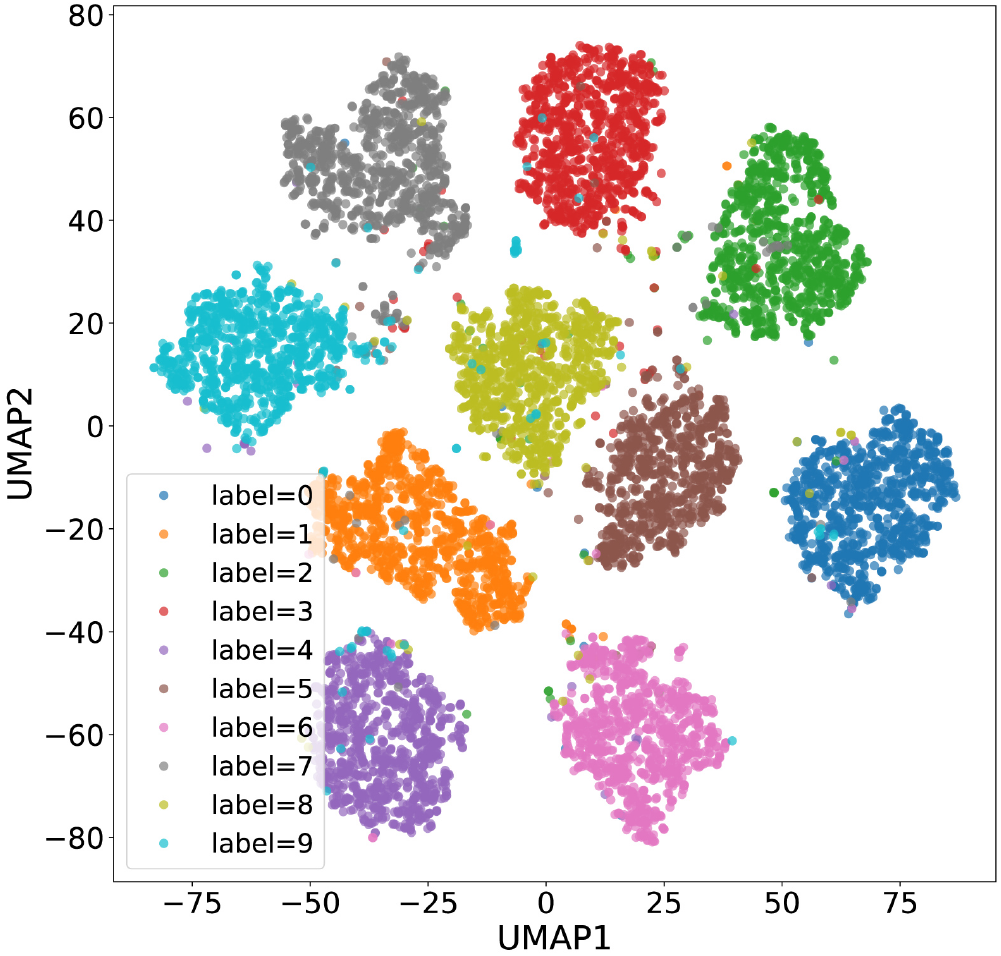
The embedding representation in the last pooling layer of the convolutional neural network exhibits well-separated cluster patterns among 10 digits on the MNIST dataset.

## Footnotes

1 The Apache licensed source code of ACE will be available at bitbucket.org/noblelab/ace.

